# Cognitive Model Discovery via Disentangled RNNs

**DOI:** 10.1101/2023.06.23.546250

**Authors:** Kevin J. Miller, Maria Eckstein, Matthew M. Botvinick, Zeb Kurth-Nelson

**Affiliations:** Google DeepMind and University College London, London, UK; Google DeepMind, London, UK

## Abstract

Computational cognitive models are a fundamental tool in behavioral neuroscience. They instantiate in software precise hypotheses about the cognitive mechanisms underlying a particular behavior. Constructing these models is typically a difficult iterative process that requires both inspiration from the literature and the creativity of an individual researcher. Here, we adopt an alternative approach to learn parsimonious cognitive models directly from data. We fit behavior data using a recurrent neural network that is penalized for carrying information forward in time, leading to sparse, interpretable representations and dynamics. When fitting synthetic behavioral data from known cognitive models, our method recovers the underlying form of those models. When fit to laboratory data from rats performing a reward learning task, our method recovers simple and interpretable models that make testable predictions about neural mechanisms.

## 1 Introduction

Fitting quantitative cognitive models to sequential behavioral data is a fundamental tool in many areas of psychology and neuroscience [44, 11, 9, 35]. These models can be viewed as mechanistic hypotheses about the cognitive processes used by the brain. Viewed in this way, they act as an explicit software instantiation of a particular cognitive hypothesis, and can be used to make sophisticated quantitative predictions about behavioral and neural data. A traditional modeling pipeline is for a human researcher to iterate over a three-step process: first to propose a candidate model structure (e.g. Q-learning), second to optimize model parameters (e.g. learning rate) with respect to a behavioral dataset, and finally to check whether the resulting model reproduces scientifically important features of the dataset. However, discovering an appropriate model structure is difficult (there are many possible structures to explore) as well as biased (the best structure may be one that the researcher has not thought of).

An alternative approach, is fitting recurrent neural networks (RNNs) directly to behavior using supervised learning [14, 17, 39, 2]. RNNs are highly expressive and can approximate a wide variety of model structures in their weights. This is therefore a way of discovering a well-fitting model from data automatically. The drawback of this approach is that the resulting RNN is a “black box”: a complex system that itself requires further analysis if it is to yield insight into cognitive mechanism.

Here, we propose a solution that aims to achieve the best of both worlds: automated discovery of cognitive models that are human-interpretable. Our approach, which we call disentangled RNN or “DisRNN”, draws on recently developed methods from machine learning for “disentangling” [21, 22, 7, 19]. These methods encourage networks to learn representations in which each dimension corresponds to a single true factor of variation in the data [21, 7]. DisRNN encourages disentangling in two ways. The first is to separate the update rule for each element of the latent state into separate sub-networks. The second is to use information “bottlenecks”, which impose a penalty on maintaining information within the network, to the inputs and outputs of these sub-networks.

We fit DisRNN on sequential behavioral datasets from rats and artificial agents performing a classic cognitive task, the dynamic two-armed bandit [12, 26, 24, 38]. First, we generate synthetic datasets using artificial agents with known learning algorithms, Q-Learning and Actor-Critic, which have different update rules and carry different information between timesteps. When fitting the behavior output of these algorithms, DisRNN correctly recovers the timecourses of latent state information and the structure of the update rules. Second, we fit DisRNN on large laboratory datasets generated by rats performing the same task [34]. We find that DisRNN provides a similar quality of fit to the best known human-derived cognitive model of this dataset [34] as well as to an unconstrained neural network [14]. We find that DisRNN learns simple human-interpretable cognitive strategies which can be used to make predictions about behavioral and neural datasets.

## 2 Related Work

Our strategy for encouraging networks to adopt disentangled representations is directly inspired by work using variational autoencoders (VAE). Specifically, *β*-VAE [21, 7], by scaling the KL loss of a variational autoencoder’s sampling step (which can be viewed as requiring information to pass through a Gaussian information bottleneck), learns sparse, disentangled representations in which each latent variable corresponds to a single true factor of variation in the data. While this work considered feedforward autoencoders, a wide variety of techniques have been proposed combining elements of feedforward VAEs with recurrent neural networks [18]. We adapt these ideas here in a way that emphasizes interpretability and is appropriate for cognitive model discovery.

While the use of recurrent neural networks as cognitive models has a long history [5, 25], recent work has expanded the toolkit for fitting networks to laboratory behavioral datasets and interpreting them [14, 17, 39, 13]. This approach of fitting standard neural networks and attempting to interpret them is complementary to our own, and benefits from the growing toolkit for neural network interpretability [40, 32, 33]. Our approach is complementary: instead of fitting standard networks and developing tools to interpret them post hoc, we develop networks which are incentivized to learn easily-interpretable solutions. Within neuroscience, several methods exist which attempt to discover interpretable latent dynamics from neural recording data [36, 43]. To our knowledge, these ideas have not been applied in the context of cognitive model discovery from behavioral data.

The two-armed bandit task we consider here is one of a family of dynamic reward learning tasks that have been heavily studied in behavioral neuroscience. This has led to a large library of candidate cognitive models [12, 4, 15, 29, 10, 24, 31, 34, 30, 3, 37]. The development of such models typically follows a theory-first approach, beginning with an idea (e.g. from optimality [12, 37], from machine learning [30], or from neurobiology [31]). Several studies have fit behavioral dataset using highly flexible classical models (Markov models and logistic regressions)[24, 34, 29, 10]. One of these, the source of the dataset we use, has attempted to use these fits as the basis for cognitive model discovery by compressing the revealed patterns into a simpler model which is cognitively interpretable [34].

Two very recent papers have proposed a very similar workflow of discovering cognitive strategies from behavioral data using constrained neural networks. The first uses “hybrid” models that combine elements of classic structured models and flexible neural networks [16]. The second constrains networks by limiting them to a very small number of hidden units [2]. Future work should compare these approaches directly on matched datasets, as well as explore other possible opportunities for using the machinery of modern machine learning for model discovery.

## 3 Disentangled Recurrent Neural Networks

An RNN trained to match a behavioral dataset containing temporal dependencies [14, 17, 39] can be viewed as maintaining a set of *latent variables* which carry information from the past that is useful for predicting the future. The weights of the network can be viewed as defining a set of *update rules* defining how each latent variable evolves over time based on external observations as well as the previous values of the other latent variables. Standard RNN architectures (such as the LSTM [23]) typically learn high-dimensional latent representations with highly entangled update rules. This makes it difficult to understand which cognitive mechanisms they have learned, limiting their usefulness as cognitive hypotheses.

### 3.1 Network Architecture

In order to learn an interpretable cognitive model, we encourage sparsity and independence by imposing bottlenecks which penalize the network for using excess information (Figure 1, left) [42, 1, 28, 21, 22, 7, 19]. The bottlenecks in our networks limit information flow using Gaussian noise. Our implementation uses multiple information bottlenecks, each of which is a noisy channel defined by two learned parameters: a “multiplier” *m* and noise variance *σ*. The output on each timestep is sampled from a Gaussian distribution:

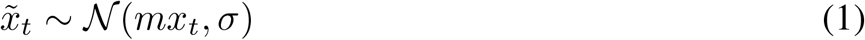

where *x* is the scalar input to the bottleneck and 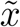 is the scalar output. Each bottleneck is associated with a loss which penalizes the sampling distribution for deviating from the unit Gaussian:

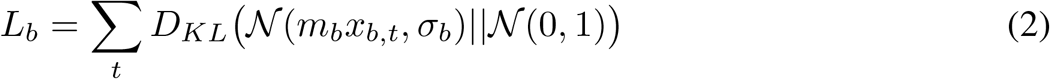

where *D*_*KL*_ is the Kullback-Leiber divergence, which quantifies the difference between the two probability distributions. This difference, and therefore the cost of the bottleneck, will be zero in the case that *m* = 0 and *σ* = 1. In that situation, the bottleneck will output samples from the unit Gaussian. These outputs will be independent of the input, meaning that no information will flow through the bottleneck. We refer to these as “closed bottlenecks”. In all other situations, *L*_*b*_ will be greater than zero, and some information will pass through. In our experiments, bottlenecks that are carrying useful information typically fit *m* ≈ 1 and *σ* ≪ 1. We refer to these as “open bottlenecks”.

**Figure 1.**
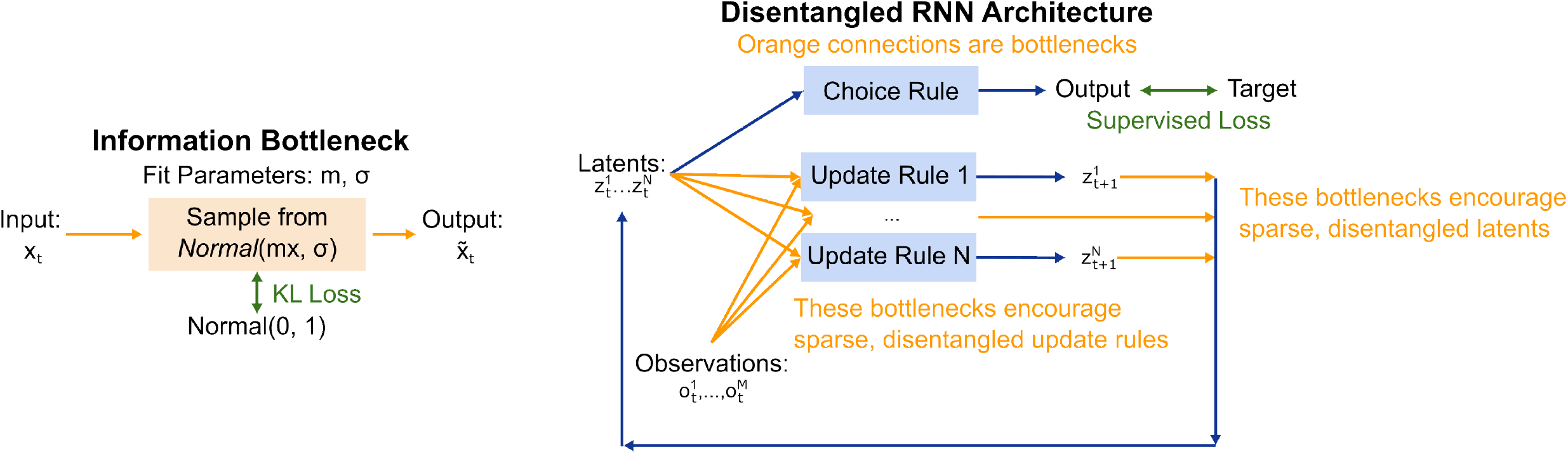
Network Architecture. **Left:** A single information bottleneck. Output 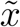 is a random sample from a Gaussian distribution determined by input *x* and bottleneck parameters *m* and σ. The bottleneck is associated with a KL loss penalizing information transfer. **Right**: Overall architecture of the DisRNN. Each latent variable is updated by a separate feedforward neural network. Bottlenecks (orange connections) are imposed both on the inputs to these networks and on each latent variable to encourage interpretable representations.

We use these information bottlenecks to encourage cognitively interpretable models by encouraging two distinct kinds of disentangling. The first is disentangling in the latent variables themselves: we would like these to capture separable latent processes and be relatively few in number. We encourage this first kind of simplicity by imposing a separate bottleneck on each scalar element of the network’s hidden state. We refer to these as the “latent bottlenecks”.

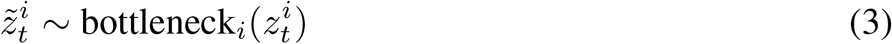

The second kind of simplicity is in the update rules for the latent variables. We would like each variable to be updated by its own separate rule, and we would like each of these rules to be as simple as possible. We encourage this kind of simplicity by updating each element of the network’s hidden state, *z*_*i,t*_, using a separate learned update rule defined by a separate set of parameters. Each update rule consists of a multilayer perceptron (MLP) which defines a multiplicative update to be applied to its corresponding latent variable. These “Update MLPs” have access to the values of all elements of the previous timestep’s hidden state, 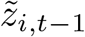 (having passed already through its latent bottleneck), as well as to the network’s current observations *o*_*t*_. Each element of the MLP input (i.e. of 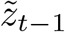 and of *o*_*t*_) must pass through an additional information bottleneck, which we refer to as “Update MLP Bottlenecks”. The output of each update MLP is a scalar weight *w* and update target *u*, which are used together to update the value of the associated latent variable *z* in a manner analogous to the Gated Recurrent Unit [8].

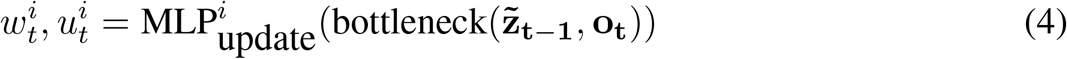

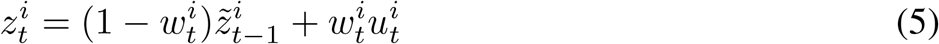

The latent variables are used on each timestep to make a prediction about the target, *y*, using a separate “Choice MLP”:

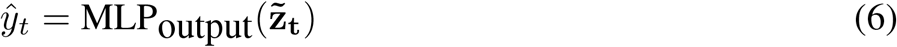

We train the model end-to-end by gradient descent to minimize the total loss function:

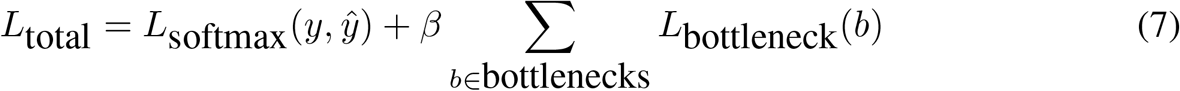

where *y* are supervised targets, *L*_softmax_ is a softmax cross-entropy loss. The hyperparameter *β* scales the cost associated with passing information through the bottleneck [21, 7]. Different values of *β* are expected to produce solutions which adopt different tradeoffs between predictive accuracy and model simplicity.

### 3.2 Training Details

All networks trained for this paper were of the same size and trained in the same way. Each network had five latent variables. Update MLPs consisted of three hidden layers containing five units each. The Choice MLP consisted of two hidden layers of two units each. We used the rectified linear (ReLU) activation function. Networks were defined using custom modules written using Jax [6] and Haiku [20]. Datasets consisted of sequences of binary choices made (left vs. right) and outcomes experienced (reward vs. no reward) by either a rat [34] or an artificial agent (Q-Learning or Leaky Actor-Critic, see below). On each timestep, the observation given to the network consisted of the choice and reward received on the corresponding trial, and the target was the choice on the subsequent trial. Network parameters were optimized using gradient descent and the Adam optimizer [27], with a learning rate of 5×10^−3^. We typically trained networks for 10^5^ steps, except that networks with very low *β* (10^−3^ or 3×10^−4^) required longer to converge and were trained for 5×10^5^ steps. Using a second-generation TPU, models required between four and fifty hours to complete this number of training steps.

## 4 DisRNN Recovers True Structure in Synthetic Datasets

We first apply DisRNN to synthetic datasets in which the process generating the data is known. We consider a task that has been the subject of intensive cognitive modeling efforts in psychology and neuroscience, the dynamic two-armed bandit task (Figure 2) [12, 26, 24, 38, 34]. In each trial of this task, the agent selects one of two available actions and then experiences a probabilistic reward. In our instantiation of the task, rewards are binary, and the reward probability conditional on selecting each arm drifts independently over time according to a bounded random walk [34].

**Figure 2.**
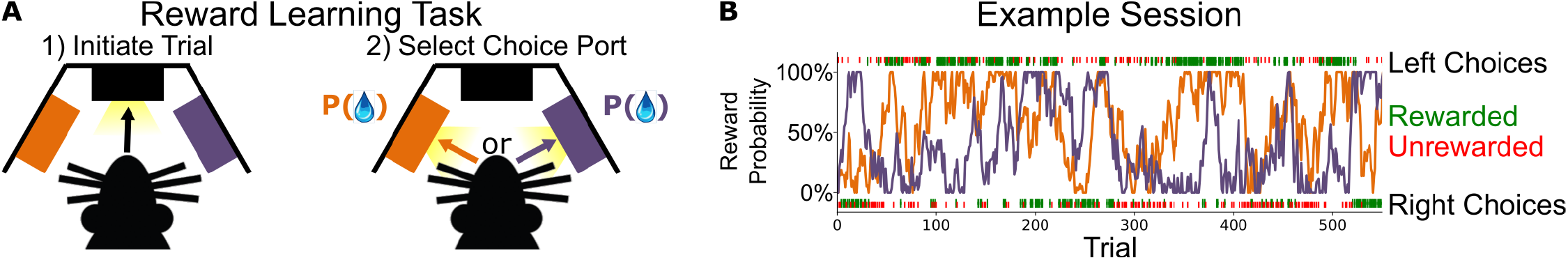
Dynamic Two-Armed Bandit Task. **a**: In each trial, the agent (rat or synthetic agent) selects one of two possible actions (left or right) and receives one of two possible outcomes (reward or no reward). **b**: Example rat behavioral session from [34]. Orange and purple lines show the drifting ground truth reward probabilities for each action. The placement of each tick mark (above or below) indicates the behavior of the animal on that trial (chose left or chose right), while the color of the tick indicates whether the animal received reward. We will use this session as a running example to illustrate the dynamics of artificial agents and RNNs.

We generated synthetic datasets from two reinforcement learning agents performing this task: Q-Learning and Leaky Actor-Critic. These agents have markedly different latent variables and update rules, allowing us to test whether DisRNN can recover the correct structure of each agent.

### 4.1 Q-Learning Agent

The Q-learning agent [41] maintains two latent variables, *Q*_left_ and *Q*_right_. Each of these is associated with one of the available actions, and gives a running average of recent rewards experienced after taking that action. They are updated according to the following rules:

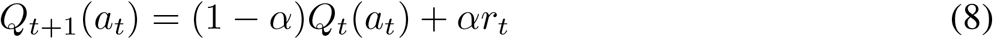

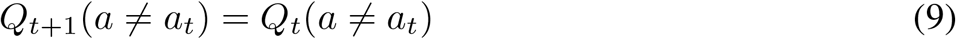

We visualize these update rules by plotting the updated value of *Q* as a function of its initial value and of the trial type (Figure 3a). A key feature is that each *Q* value only updates when its corresponding action is selected.

**Figure 3.**
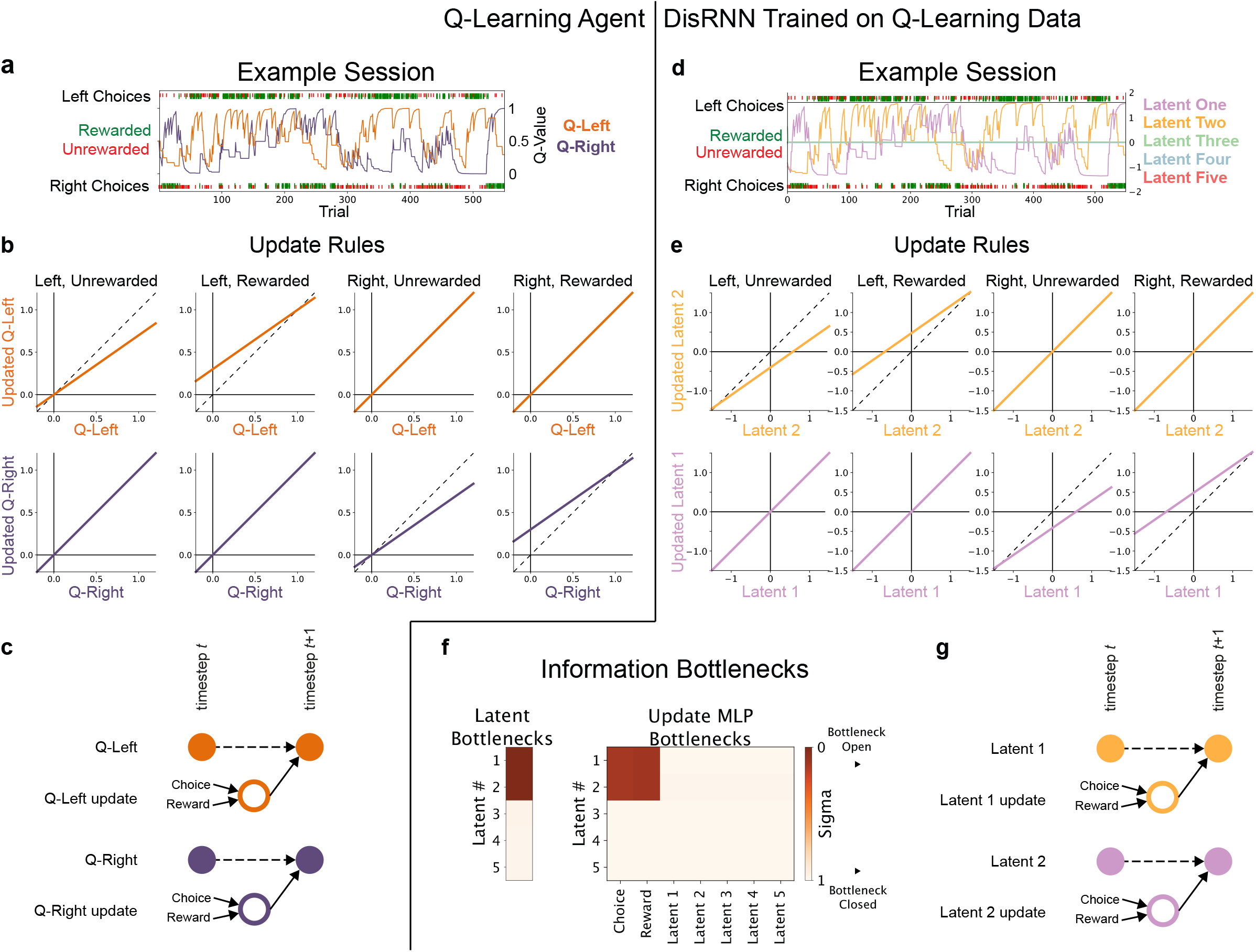
DisRNN Recovers Latent Dynamics of Q-Learning. **a**: Q-Learning agent run using the choices and rewards from the example behavioral session (Figure 2) to generate timeseries for *Q*_left_ and *Q*_right_. **b**: Visualization of the Q-Learning update rules. Each panel visualizes the update of either *Q*left (top row) or *Q*right (bottom row) following a particular combination of choice and outcome (columns). The post-update value is shown on the vertical axis, as a function of the pre-update value, on the horizontal. Dashed lines are identity. **c**: Dependency graph of Q-learning. Q-learning computes an update for each latent (*Q*_left_ and *Q*right) on each timestep. The update for *Q*_left_ depends on choice and reward (Equation 8). The new value of *Q*_left_ is a weighted sum of its old value (dashed line) and the update. *Q*right is similarly updated. **d**: DisRNN trained on a synthetic behavioral dataset generated by the Q-Learning agent. This panel shows data from the trained model run with choice and reward input from the same example session. Latents have been ordered, signed, and colored to highlight similarities with the Q-Learning agent. **e**: Visualization of the learned update rules (equations 4 and 5) for the first two latents. **f**: Learned parameters of the information bottlenecks. Left: Transmission bottlenecks (equation 3). Right: Update MLP bottlenecks (equation 4). Darker colors indicate open bottlenecks: for example, the update for latent 1 can depend on choice and reward. **g**: Dependency graph of cognitive model learned by DisRNN. Note that this graph was obtained automatically from the data, and exactly matches the graph used to generate the data.

The Q-Learning agent selects actions based on the difference in *Q*-values:

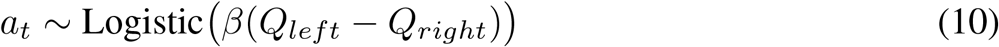

We generated a large synthetic dataset consisting of choices made and rewards received by the Q-Learning agent (*α* = 0.3, *β* = 3; 1000 sessions of 500 trials each) (Figure 3b). We then trained a DisRNN to imitate this dataset. Inspecting the learned bottleneck parameters of this network (Figure 3e), we find that just two of its latents have open bottlenecks (*sigma*≪ 1), allowing them to pass information forward through time. We find that the update rules for these latents have open bottlenecks for the input from both previous choice and previous reward. We find that the three latents with closed (*σ*≈1) bottlenecks take on near-zero values on all timesteps, while the two with open bottlenecks follow timecourses that are strikingly similar to those of *Q*_*left*_ and *Q*_*right*_ (albeit centered on zero instead of on 0.5; Figure 3c). We visualize the learned update rule associated with each latent by probing its associated update MLP (Figure 3d). We find that these update rules are strikingly similar to those for *Q*_*left*_ and *Q*_*right*_. They reproduce the key feature that each is only updated following choices to a particular action.

### 4.2 Leaky Actor-Critic Agent

The Leaky Actor-Critic agent uses a modification of the policy gradient learning rule [41](Section 2.8). Like the Q-Learning agent, it makes use of two latent variables, but uses very different rules to update them. The first variable tracks “value”, *V*. It keeps a running average of recent rewards, regardless of the choice that preceded them:

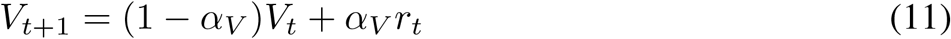

where *α*_*V*_ is a learning rate parameter.

The second is a “policy” variable Θ, which determines the probability with which the agent will select each action. The policy gradient algorithm defines two variables *θ*_*left*_ and *θ*_*right*_, each associated with one of the actions and updated using the following rules:

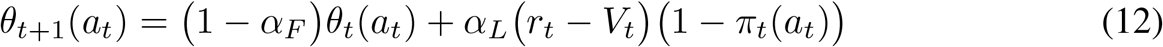

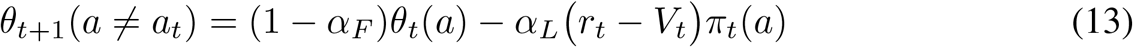

where *π*_*t*_ is the agent’s policy defined below (equation 14), *α*_*L*_ is a learning rate parameter, and *α*_*F*_ is a forgetting rate parameter that we have added to the usual update rule. This forgetting mechanism causes the policy variables to gradually decay towards zero over time, allowing the agent to perform well in an environment with constantly-changing reward probabilities. In a setting with just two available actions, the above update rules cause *θ*_left_ and *θ*_right_ to be degenerate: it will always be the case that *θ* _left_= −*θ* _right_. We therefore define a single latent variable Θ = *θ* _left_− *θ* _right_ for the purposes of visualizing this agent. The Leaky Actor-Critic agent selects actions according to:

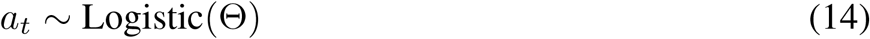

We visualize these update rules by plotting the updated values of *V* and of Θ as a function of their initial value and of the trial type (Figure 4b). These update rules are quite different from those of the Q-Learning agent (Figure 3). One key feature is that the update rule for *V* depends only on reward, not on choice. Another key feature is that the update rule for Θ depends not only on choice and reward, but also on the current value of *V*.

**Figure 4.**
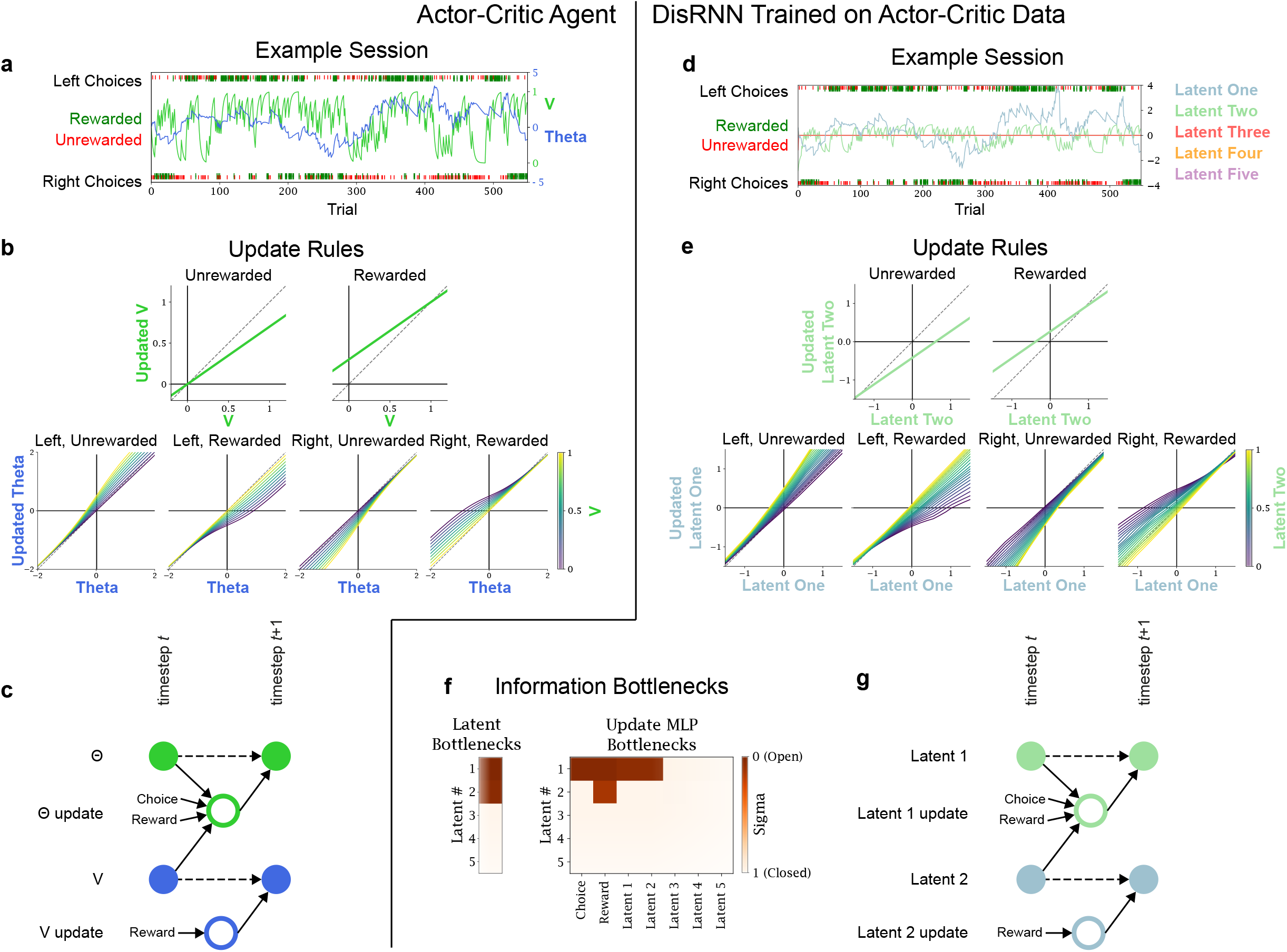
DisRNN Recovers Latent Dynamics of Leaky Actor-Critic. **a**: Actor-Critic agent run on the example rat behavioral session (Figure 2) to generate timeseries for *V* and Θ. **b**: Visualization of the Actor-Critic update equations. The update for *v* depends on reward but is independent of choice. The update for Θ depends on choice and reward, as well as the value of *V*. **c**: Dependency graph of Actor-Critic learning. **d**: DisRNN trained on a synthetic behavioral dataset generated by an Actor-Critic agent. **e**: Visualization of the learned update rules for the first two latents. **f**: Learned bottleneck parameters. Left: Transmission bottlenecks (equation 3). The first two latents have open bottlenecks; the remaining three latents have closed bottlenecks. Right: Update MLP bottlenecks (equation 4). **g**: Dependency graph of DisRNN trained on Actor-Critic data.

We generated a large synthetic dataset consisting of choices made and rewards received by the Leaky Actor-Critic agent (*α*_*V*_ = 0.3, *α*_*L*_ = 1, *α*_*F*_ = 0.05; 1000 sessions of 500 trials each), and trained a DisRNN to imitate this dataset. Inspecting the learned bottleneck parameters of this network (Figure 4e), we find that two of its latents have open bottlenecks allowing them to pass information forward through time. One of these (latent two) has an open bottleneck to receive input from reward only. We find that this latent is strikingly similar to *V* from the Leaky Actor-Critic agent, both in terms of its timecourse (Figure 4c) and update rule (Figure 4d, above). The other (latent one) has open bottlenecks to receive input from choice, reward, and from latent two. This latent is strikingly similar to Θ from the Leaky Actor-Critic Agent, both in terms of its timecourse (Figure 4c) and update rule (Figure 4d, below).

Together with the previous section, these results indicate that DisRNN is capable of recovering the true generative structure of agents with two very different cognitive strategies using behavioral data alone.

## 5 DisRNN Reveals Interpretable Models of Rat Behavior

Having established that DisRNN can recover true cognitive mechanisms in synthetic datasets where ground truth is known, we moved on to consider a large laboratory dataset from rats performing the drifting two-armed bandit task [34]. This dataset has previously been the subject of an intensive human effort at data-driven cognitive modeling, using an approach of successive model reduction. This approach resulted in a particular cognitive model consisting of three components each with its own latent variable: a fast-timescale reward-seeking component, a slower perseverative component, and a very slow “gambler’s fallacy” component. This model provided a better fit to the dataset than existing models from the literature, and is the best known cognitive model for this dataset [34]. We first asked whether DisRNN could provide a similar quality of fit to this model. We fit DisRNNs using four different values of the parameter *β* (10^−3^, 3×10^−3^, 10^−2^, and 3×10^−2^) that controls the relative weight of predictive power and model simplicity in the network’s loss function (equation 7). Following [34], we evaluated model performance using two-fold cross-validation: we divide each rat’s dataset into even-numbered and odd-numbered sessions, fit a set of model parameters to each, and compute the log-likelihood for each parameter set using the unseen portion of the dataset. To compare results across different animals, we use “normalized likelihood”: *e*log-likelihood*/*n trials [11]. For each rat, we compute this cross-validated normalized likelihood both for our DisRNNs and for the cognitive model from [34], and plot the differences in Figure 5. We find that DisRNNs trained with *β* = 3×10^−3^ achieved a quality of fit similar to the cognitive model, and that those with *β* = 10^−3^ even narrowly outperformed it (Figure 5, orange curve). As an additional benchmark, we also compared our models to a widely-used RNN architecture, the LSTM [23]. We selected LSTM hyperparameters (network size and early stopping) using subject-level cross-validation: we divided the dataset into subsets containing only even-numbered and odd-numbered rats, identified the hyperparameters that maximized (session-level) cross-validated likelihood in each subset, and evaluated networks with those hyperparameters in the unseen subset. We found that DisRNNs with *β* = 10^−3^ achieved a very similar quality of fit to these optimized LSTMs. Together these results indicate the DisRNNs, despite their architectural constraints and disentanglement loss, can achieve predictive performance comparable to standard RNNs and to well-fit human-derived cognitive models.

**Figure 5.**
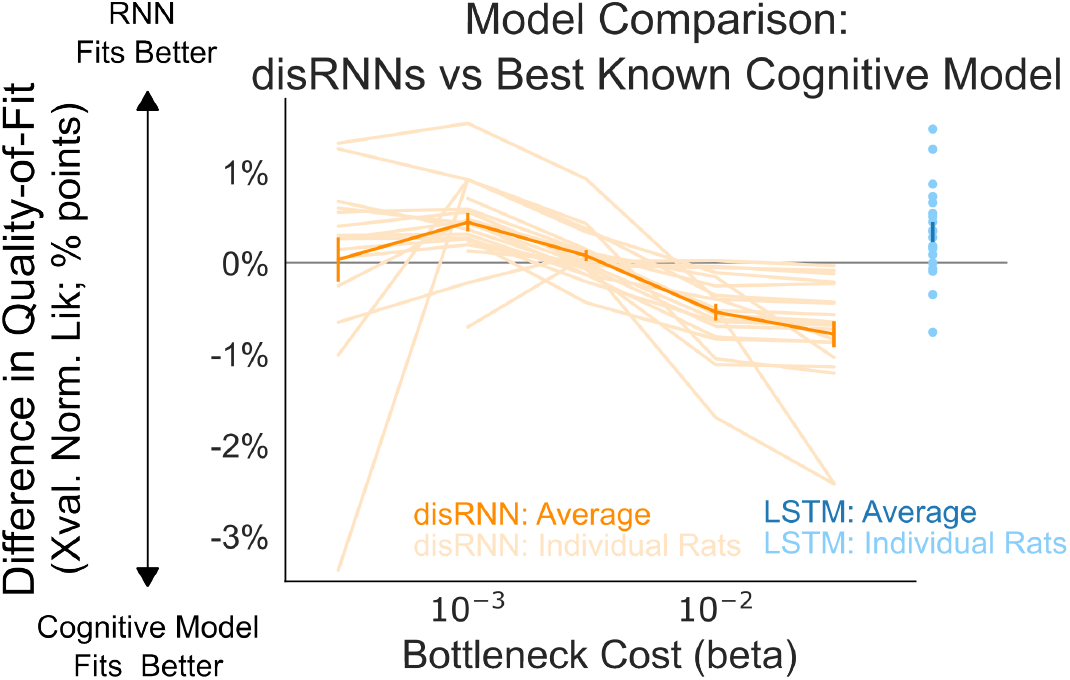
Model Comparison on Rat Behavioral Dataset. The y-axis shows performance (cross-validated normalized likelihood) relative to the human derived cognitive model from [34]. DisRNN is in orange, and LSTM is in blue. The x-axis for DisRNN represents different values of the parameter *β*, which controls the tradeoff between model simplicity and predictive performance. LSTM hyperparameters were selected by subject-level cross-validation. *N* = 20 rats in light traces, means in heavy traces; error bars indicate standard error over rats.

Finally, we examined the parameters of DisRNNs fit to the complete dataset for each rat. We found that these were typically low-dimensional, with only a small number of latent variables having open bottlenecks, as well as sparse, with each update MLP having only a small number of open bottlenecks. While a thorough characterization of the patterns present in DisRNNs fit to individual rats remains a direction for future work, we present a representative example in Figure 6. This figure shows three DisRNNs with different values of *β* fit to the dataset from the same example rat. The DisRNN with the largest *β* of 3×10^−2^ (left) adopts a strategy that utilizes only one latent variable. This strategy mixes reward-seeking and perseverative patterns into a single update rule (note that fixed points, where the line crosses unity, are more extreme for the rewarded conditions than the unrewarded ones). The DisRNN with a medium *β* of 10^−2^ (middle) adds a second latent variable that is purely perseverative and operates on a considerably slower timescale. Its fast-timescale reward-seeking latent variable, however, shows a similar perseverative signature to the previous network. The network with the smallest *β* of 3×10^−3^ (right) retains both the fast reward-seeking and slower perseverative dynamics and adds a third, much slower-timescale, latent. The update rule for this latent depends both on reward and on the value of the perseverative latent. Interestingly, this very slow latent influences the update rules for both the reward-seeking and the perseverative latents. While the best human-derived cognitive model does include a very long-timescale component, this component simply influences choice rather than modulating the update rules of the other components. Situations where one latent variable modulates the update rule of another do sometimes occur in human-derived models for related tasks [3, 37], though it is not clear whether the pattern seen in this network is consistent with any of these. Characterizing these patterns in more detail, and especially quantifying similarities and differences among different animals, will be an important step for future work.

**Figure 6.**
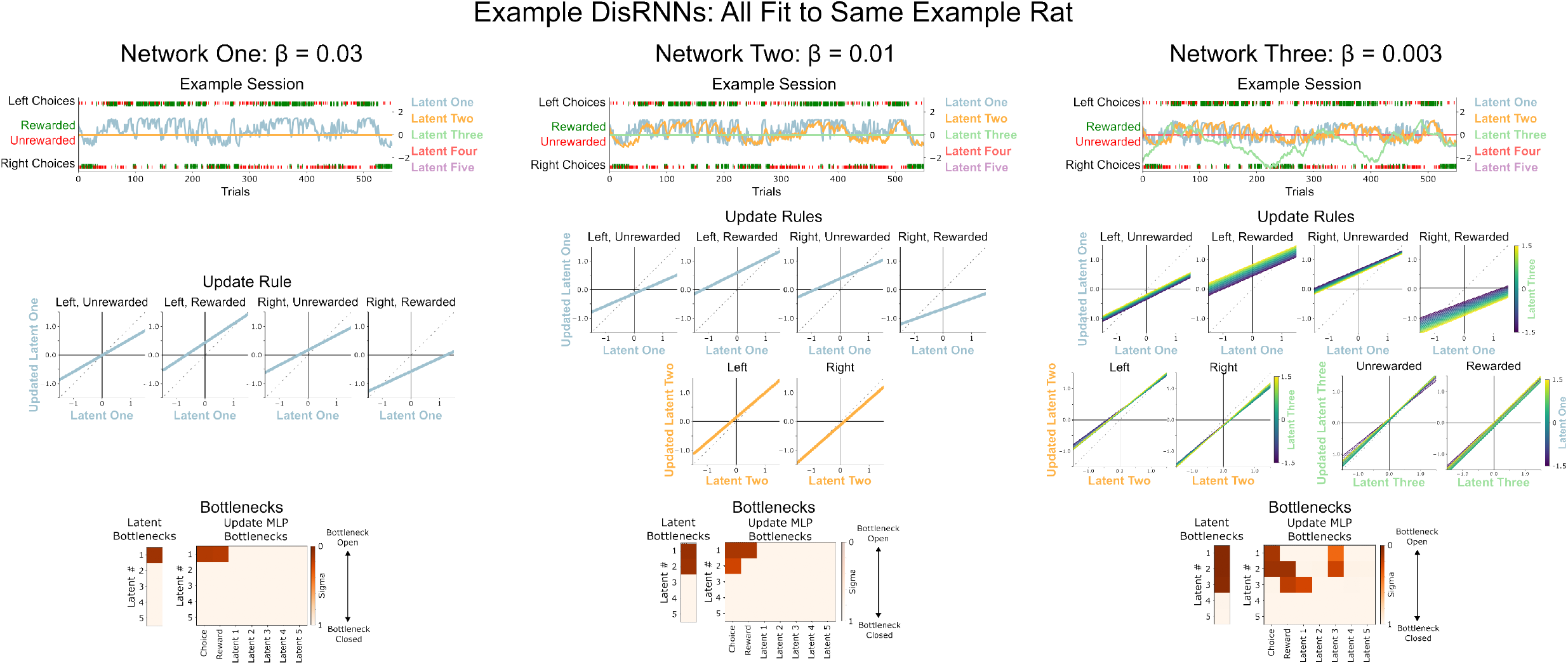
DisRNN Trained on Rat Datasets. Examples of fit DisRNN networks with different values of the hyperparameter *β* controlling the tradeoff between simplicity and predictive power.

## 6 Discussion

In this work, we develop a framework for discovering parsimonious cognitive models directly from behavioral data by fitting recurrent neural networks that contain several structural features that encourage them to learn sparse, disentangled representations. Fit to synthetic datasets generated using known cognitive mechanisms, our method accurately recovers the structure of those mechanisms. Fit to laboratory datasets from rats performing a cognitive task, our method reveals models that are relatively simple and human-interpretable models while outperforming the best known human-derived cognitive model in terms of predictive accuracy.

The fit disRNNs have several key features than enable them to be applied immediately to standard cognitive neuroscience workflows [44, 11, 9, 35]. The first is that they generate timestep-by-timestep timecourses for the values of latent variables that play known cognitive roles within the model. These timecourses can be used as predictions about neural activity: if the model’s mechanisms are implemented in the brain, then somewhere in the brain there is likely to be a signal that follows a similar timecourse. A second key feature is that disRNN makes explicit the rules by which latent variables are updated. These can be used as predictions about update rules within the brain: if the model’s mechanisms are implemented in the brain, then somewhere in the brain there are likely to be similar update rules (e.g. synaptic weights to update the ongoing neural activity, synaptic plasticity to adjust synaptic weights, etc). A third is that a trained disRNN can be used to generate predictions for experiments which alter neural activity, for example by silencing it using optogenetics. Activity within the disRNN can be altered, for example by artificially zero-ing out a particular latent on a particular subset of trials, and the model can be run to generate predictions for behavioral and neural data.

The most important limitation of our approach is that there is no guarantee that the recovered model will necessarily correspond to the true cognitive mechanisms used by the brain. Instead, it uses the dataset to reveal a parsimonious hypothesis. This hypothesis that requires evaluation by a human scientist to determine whether, given the rest of what is known about the psychology and biology of the system, it is plausible. If plausible, it may require new experiments to test the predictions that it makes. This limitation is fundamental to any approach that seeks to make inferences about cognitive mechanisms by analyzing behavioral data, as any behavioral dataset will provide only a limited window on neural mechanisms. One view of these systems is that they perform automatic hypothesis generation, without themselves addressing the problem of hypothesis testing.

Other limitations are that the method, because it requires fitting quite flexible models, likely is applicable only to relatively large-scale datasets. Determining how performance scales with dataset size and exploring methods of improving data efficiency may be important directions for further research. Another direction is exploring the performance of disRNNs on behavioral datasets from other types of cognitive neuroscience tasks.

## References

[1] Alexander A Alemi et al. “Deep variational information bottleneck”. In: arXiv preprint arXiv:1612.00410 (2016).

[2] Li Ji-An, Marcus K Benna, and Marcelo G Mattar. “Automatic Discovery of Cognitive Strategies with Tiny Recurrent Neural Networks”. In: bioRxiv (2023), pp. 2023–04.

[3] Timothy EJ Behrens et al. “Learning the value of information in an uncertain world”. In: Nature neuroscience 10.9 (2007), pp. 1214–1221.

[4] Celia C Beron et al. “Mice exhibit stochastic and efficient action switching during probabilistic decision making”. In: Proceedings of the National Academy of Sciences 119.15 (2022), e2113961119.

[5] Matthew Botvinick and David C Plaut. “Doing without schema hierarchies: a recurrent connectionist approach to normal and impaired routine sequential action.” In: Psychological review 111.2 (2004), p. 395.

[6] James Bradbury et al. JAX: composable transformations of Python+NumPy programs. Version 0.3.13. 2018. URL: http://github.com/google/jax.

[7] Christopher P Burgess et al. “Understanding disentangling in beta-VAE”. In: arXiv preprint arXiv:1804.03599 (2018).

[8] Kyunghyun Cho et al. “On the properties of neural machine translation: Encoder-decoder approaches”. In: arXiv preprint arXiv:1409.1259 (2014).

[9] Greg Corrado and Kenji Doya. “Understanding neural coding through the model-based analysis of decision making”. In: Journal of Neuroscience 27.31 (2007), pp. 8178–8180.

[10] Greg S Corrado et al. “Linear-nonlinear-Poisson models of primate choice dynamics”. In: Journal of the experimental analysis of behavior 84.3 (2005), pp. 581–617.

[11] Nathaniel D Daw et al. “Trial-by-trial data analysis using computational models”. In: Decision making, affect, and learning: Attention and performance XXIII 23.1 (2011).

[12] Nathaniel D Daw et al. “Cortical substrates for exploratory decisions in humans”. In: Nature 441.7095 (2006), pp. 876–879.

[13] Amir Dezfouli, Richard Nock, and Peter Dayan. “Adversarial vulnerabilities of human decisionmaking”. In: Proceedings of the National Academy of Sciences 117.46 (2020), pp. 29221–29228.

[14] Amir Dezfouli et al. “Models that learn how humans learn: The case of decision-making and its disorders”. In: PLoS computational biology 15.6 (2019), e1006903.

[15] R Becket Ebitz, Eddy Albarran, and Tirin Moore. “Exploration disrupts choice-predictive signals and alters dynamics in prefrontal cortex”. In: Neuron 97.2 (2018), pp. 450–461.

[16] Maria K. Eckstein et al. “Predictive and Interpretable: Combining Artificial Neural Networks and Classic Cognitive Models to Understand Human Learning and Decision Making”. In: bioRxiv (2023). DOI: 10.1101/2023.05.17.541226. eprint: https://www.biorxiv.org/content/early/2023/05/17/2023.05.17.541226.full.pdf. URL: https://www.biorxiv.org/content/early/2023/05/17/2023.05.17.541226.

[17] Matan Fintz, Margarita Osadchy, and Uri Hertz. “Using deep learning to predict human decisions and using cognitive models to explain deep learning models”. In: Scientific reports 12.1 (2022), p. 4736.

[18] Laurent Girin et al. “Dynamical Variational Autoencoders: A Comprehensive Review”. In: CoRR abs/2008.12595 (2020). arXiv: 2008.12595. URL: https://arxiv.org/abs/2008.12595.

[19] Anirudh Goyal et al. “Infobot: Transfer and exploration via the information bottleneck”. In: arXiv preprint arXiv:1901.10902 (2019).

[20] Tom Hennigan et al. Haiku: Sonnet for JAX. Version 0.0.9. 2020. URL: http://github.com/deepmind/dm-haiku.

[21] Irina Higgins et al. “Beta-VAE: Learning basic visual concepts with a constrained variational framework”. In: International conference on learning representations. 2017.

[22] Irina Higgins et al. “Darla: Improving zero-shot transfer in reinforcement learning”. In: International Conference on Machine Learning. PMLR. 2017, pp. 1480–1490.

[23] Sepp Hochreiter and Jürgen Schmidhuber. “Long short-term memory”. In: Neural computation 9.8 (1997), pp. 1735–1780.

[24] Makoto Ito and Kenji Doya. “Validation of decision-making models and analysis of decision variables in the rat basal ganglia”. In: Journal of Neuroscience 29.31 (2009), pp. 9861–9874.

[25] MI Jordan. Serial order: a parallel distributed processing approach. technical report, june 1985-march 1986. Tech. rep. California Univ., San Diego, La Jolla (USA). Inst. for Cognitive Science, 1986.

[26] Hoseok Kim et al. “Role of striatum in updating values of chosen actions”. In: Journal of neuroscience 29.47 (2009), pp. 14701–14712.

[27] Diederik P Kingma and Jimmy Ba. “Adam: A method for stochastic optimization”. In: arXiv preprint arXiv:1412.6980 (2014).

[28] Diederik P Kingma and Max Welling. “Auto-encoding variational bayes”. In: arXiv preprint arXiv:1312.6114 (2013).

[29] Brian Lau and Paul W Glimcher. “Dynamic response-by-response models of matching behavior in rhesus monkeys”. In: Journal of the experimental analysis of behavior 84.3 (2005), pp. 5551–579.

[30] Daeyeol Lee, Benjamin P McGreevy, and Dominic J Barraclough. “Learning and decision making in monkeys during a rock–paper–scissors game”. In: Cognitive Brain Research 25.2 (2005), pp. 416–430.

[31] Germain Lefebvre et al. “Behavioural and neural characterization of optimistic reinforcement learning”. In: Nature Human Behaviour 1.4 (2017), p. 0067.

[32] Niru Maheswaranathan et al. “Universality and individuality in neural dynamics across large populations of recurrent networks”. In: Advances in neural information processing systems 32 (2019).

[33] Vladimir Mikulik et al. “Meta-trained agents implement bayes-optimal agents”. In: Advances in neural information processing systems 33 (2020), pp. 18691–18703.

[34] Kevin J Miller, Matthew M Botvinick, and Carlos D Brody. “From predictive models to cognitive models: Separable behavioral processes underlying reward learning in the rat”. In: bioRxiv (2018).

[35] John P O’Doherty, Alan Hampton, and Hackjin Kim. “Model-based fMRI and its application to reward learning and decision making”. In: Annals of the New York Academy of sciences 1104.1 (2007), pp. 35–53.

[36] Chethan Pandarinath et al. “Inferring single-trial neural population dynamics using sequential auto-encoders”. In: Nature methods 15.10 (2018), pp. 805–815.

[37] Payam Piray and Nathaniel D Daw. “A model for learning based on the joint estimation of stochasticity and volatility”. In: Nature communications 12.1 (2021), p. 6587.

[38] Kazuyuki Samejima et al. “Representation of action-specific reward values in the striatum”. In: Science 310.5752 (2005), pp. 1337–1340.

[39] Mingyu Song, Yael Niv, and Mingbo Cai. “Using Recurrent Neural Networks to Understand Human Reward Learning”. In: Proceedings of the Annual Meeting of the Cognitive Science Society. Vol. 43. 43. 2021.

[40] David Sussillo and Omri Barak. “Opening the black box: low-dimensional dynamics in highdimensional recurrent neural networks”. In: Neural computation 25.3 (2013), pp. 626–649.

[41] Richard S Sutton and Andrew G Barto. Reinforcement learning: An introduction. MIT press, 2018.

[42] Naftali Tishby, Fernando C Pereira, and William Bialek. “The information bottleneck method”. In: arXiv preprint physics/0004057 (2000).

[43] Alex H Williams et al. “Unsupervised discovery of demixed, low-dimensional neural dynamics across multiple timescales through tensor component analysis”. In: Neuron 98.6 (2018), pp. 1099–1115.

[44] Robert C Wilson and Anne GE Collins. “Ten simple rules for the computational modeling of behavioral data”. In: eLife 8 (Nov. 2019), e49547. ISSN: 2050-084X. DOI: 10.7554/eLife49547. URL: https://doi.org/10.7554/eLife.49547 (visited on 04/23/2021).

